# Dynamic Nucleosome Movement Provides Structural Information of Topological Chromatin Domains in Living Human Cells

**DOI:** 10.1101/059147

**Authors:** Soya Shinkai, Tadasu Nozaki, Kazuhiro Maeshima, Yuichi Togashi

**Affiliations:** Research Center for the Mathematics on Chromatin Live Dynamics (RcMcD), Hiroshima University, Higashi-Hiroshima, Japan; Biological Macromolecules Laboratory, Structural Biology Center, National Institute of Genetics, Mishima, Japan; Institute for Advanced Bioscience, Keio University, Fujisawa, Japan; Department of Genetics, School of Life Science, Graduate University for Advanced Studies (Sokendai), Mishima, Japan

## Abstract

The mammalian genome is organized into submegabase-sized chromatin domains (CDs) including topologically associating domains, which have been identified using chromosome conformation capture-based methods. Single-nucleosome imaging in living mammalian cells has revealed subdiffusively dynamic nucleosome movement. It is unclear how single nucleosomes within CDs fluctuate and how the CD structure reflects the nucleosome movement. Here, we present a polymer model wherein CDs are characterized by fractal dimensions and the nucleosome fibers fluctuate in a viscoelastic medium with memory. We analytically show that the mean-squared displacement (MSD) of nucleosome fluctuations within CDs is subdiffusive. The diffusion coefficient and the subdiffusive exponent depend on the structural information of CDs. This analytical result enabled us to extract information from the single-nucleosome imaging data for HeLa cells. Our observation that the MSD is lower at the nuclear periphery region than the interior region indicates that CDs in the heterochromatin-rich nuclear periphery region are more compact than those in the euchromatin-rich interior region with respect to the fractal dimensions as well as the size. Finally, we evaluated that the average size of CDs is in the range of 100-500 nm and that the relaxation time of nucleosome movement within CDs is a few seconds. Our results provide physical and dynamic insights into the genome architecture in living cells.

**Author Summary:** The mammalian genome is partitioned into topological chromatin domains (CDs) in the living cell nuclei. Gene expression is highly regulated within CDs according to their structure, whereas chromatin itself is highly dynamic. This raises the following question: how is the CD structure in such dynamic chromatin? We developed a conceptual framework that unifies chromatin dynamics and structure. Using a polymer model with a fractal domain structure in a viscoelastic medium, we analytically show that nucleosome movement is subdiffusive and depends on CD structure. Hence, structural information can be extracted based on nucleosome movement in living cells with single-particle tracking experiments. This framework provides physical insights into the relationship between dynamic genome organization and gene expression.

## Introduction

Genomic DNA is packed and folded three-dimensionally in the cell nuclei. In the nuclei of eukaryotic cells, the nucleosome is a basic unit consisting of an approximately 147-bp DNA wrapped around core histones [1]. Recent experimental evidences suggest that the nucleosome is irregularly folded without the 30-nm chromatin fiber [2–7]. On the other hand, at the scale of the whole nucleus, interphase chromosomes occupy distinct chromosome territories [8]. This highly organized chromosome structure allows for effective regulation of various genome functions.

By virtue of recent developments of chromosome conformation capture (3C) techniques, the genome-wide chromosome organization has been revealed by detecting the physical contact frequencies between pairs of genomic loci [9]. More recently, 3C derivatives, Hi-C and 5C profiles demonstrated that metazoan genomes are partitioned into submegabase-sized chromatin domains (CDs) including topologically associating domains (TADs) [10–12]. TADs are considered to be a regulatory and structural unit of the genome [13]; genome loci located in the same TAD are associated with each other, whereas genomic interactions are sharply depleted between adjacent domains. For even single-cell Hi-C, individual chromosomes maintain domain organization [14]. Furthermore, kilobase-resolution in situ Hi-C maps identified not only small contact domains but also CTCF-mediated loop domains [15,16].

In contrast, dynamic aspects of chromatin have been shown by live-cell imaging experiments [17–24]. In particular, single-nucleosome imaging in living mammalian cells has revealed local nucleosome fluctuations caused by the thermal random force [25–27]. The mean-squared displacement (MSD) of dynamic nucleosome movement clearly shows subdiffusive motion, 
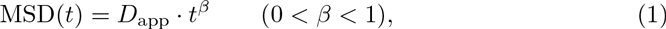
where *D_app_* is the apparent diffusion coefficient with dimension m^2^/s^β^. This means that nucleosome movement must be affected by restrictions from some factors but thermal noise. Therefore, there must be a way that the dynamic aspect is consistent with aspects of the genome organization. A theory is required to relate the dynamic aspects described by *D_app_* and *β* to the structural features of CDs. To date, the subdiffusive exponent *β* has been considered to depend on the folding structure of nucleosome fibers [28] and the viscoelasticity of the thermal environment [29,30].

The fractal nature of chromatin architecture as well as nucleus environment has been revealed recently [9,31,32]. The topological structure of CDs can be described by use of the fractal manner. Here, we propose a polymer model for a CD, whose conformational state is assumed to be expressed by the fractal dimension *d_f_* in a viscoelastic medium with the exponent 0 < α <1. Although not only the strings and binders switch model [33] but also the block copolymer model [34] can explain aspects of chromatin folding and chromosome architecture in Hi-C experiment datasets, in our model we abstract information on the conformational states of CDs and interpret their dynamic features by using size scaling according to the fractal dimensions. Accordingly, the analytical form of the MSD of nucleosomes in CDs can be derived in terms of polymer physics. As a result, the structural information of CDs, such as the size and conformational state expressed by the fractal dimension, can be derived from the MSD data of dynamic nucleosomes.

## Results

### Polymer model

#### CDs characterized by fractal dimensions

To construct a model of CDs, we assumed that a nucleosome fiber is represented as a polymer bead chain and forms a CD with size scaling, ⟨*R*⟩_CD_ ~ *N*^1/*d*_f_^ (Fig 1A), where *N* is the number of nucleosome beads in the CD, and ⟨·⟩_CD_ represents the average for all nucleosome beads within the CD at thermal equilibrium. In polymer physics, the exponent 1/*d*_f_ corresponds to the size exponent *ν* [35,36]. A nucleosome fiber in a CD not only has the excluded volume as a physical polymer, but also forms chromatin loops for transcriptional regulation [15,16,37]. Therefore, nucleosome fibers can interact with each other within the same CD through both attractive and repulsive interactions. Here, we assume that the effective conformational state of CDs is phenomenologically represented by the fractal dimension. Note that the states with df = 1, 2, and 3 correspond to a straight line, the ideal chain [35], and the fractal globule [9,28,38], respectively (Fig 1A).

**Fig 1.**
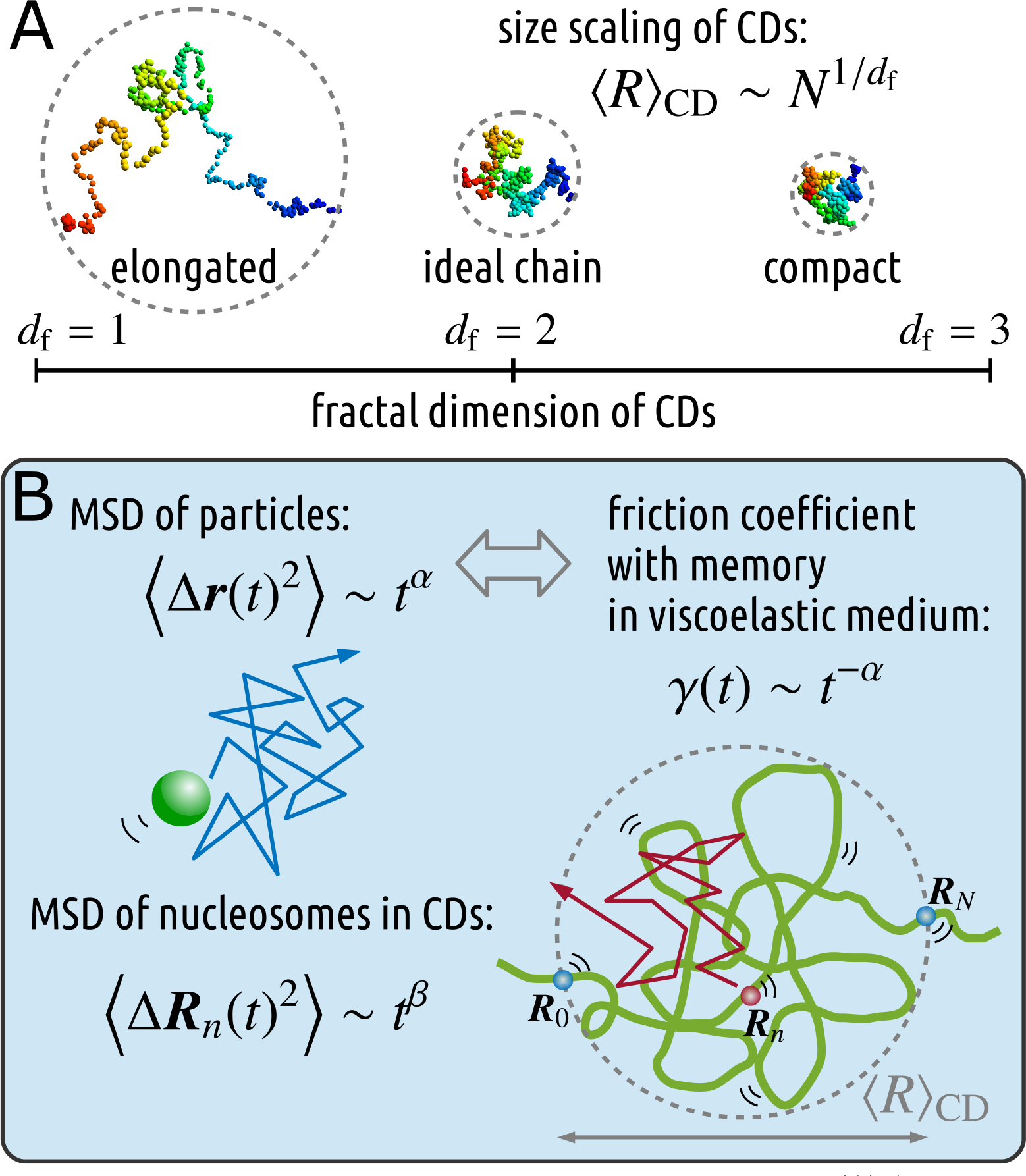
Schematic illustration of our polymer model for CDs. (A) A nucleosome fiber is represented as a polymer bead chain and forms a CD. The size scaling of CDs is expressed as *⟨R⟩_CD_ ~ N^1/df^*, where the fractal dimension represents the effective conformational state of CDs: *d_f_* = 1, 2, and 3 correspond to a straight, line, the ideal, chain, and the fractal, globule, respectively. (B) The viscoelasticity of the, medium, where the movement of particles shows the subdiffusive FBM *⟨Δr*^(t^)^2⟨^ *~ t^α^*, is described using the friction coefficient with, memory, *γ (t) ~ t ^α^*. When a nucleosome with coordinates ***R**_n_ (t)* dynamically fluctuates in the viscoelastic, medium, the movement of nucleosomes in CDs shows subdiffusion: *⟨Δ**R**_n_*(t)^2^*⟩* ~ *t^β^*

#### Nucleosome fiber fluctuation in viscoelastic medium with memory

The subdiffusive motion of tracer particles in living cells, ⟨[**r**(t) — **r**(0)⟩^2^) ~ *t^α^,* has been observed [29,39–41]. There are several physical models for generating subdiffusion, including: (i) the generalized Langevin equation (GLE), which is consistent with fractional Brownian motion (FBM) [42–45], and (ii) the continuous-time random walk [46]. Since some experiments have shown that the movement of chromosomal loci displays the FBM [23,29], here, we adopt the former model to describe the friction effect with memory in the viscoelastic medium [39,47,48] that satisfies the fluctuation-dissipation relation (FDR) [35,49]: the GLE 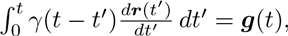, with friction coefficient with memory of γ(*t*) ~ *t^-α^,* generates the subdiffusive FBM. The thermal random force **g**(*t*) satisfies the FDR ⟨g_K_(t)g_λ_(t′)⟩ = *k_B_T_γ_(t — t′)δ_K_λ*, where *k_B_* is the Boltzmann constant, *T* is the temperature of the environment, and the suffixes *κ* and λ represent *x, y* and z.

Here, we focus on the concrete description of our polymer model. A CD is assumed to be formed by *N* +1 nucleosome beads at positions {***R***_0_, ***R***_1_, …, ***R***_N_} (Fig 1B), and adjacent beads are connected via a harmonic spring so that the effective bond length is *b*_eff_, and long-range interactions exist such that the phenomenological size scaling of the CDs is proportional to *N^1/d_f_^*. Moreover, as mentioned above, the friction effect between each nucleosome and the viscoelastic medium is assumed to be described by the friction coefficient with memory [30,44,45,47],

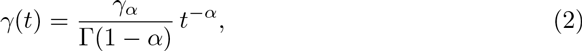
where the dimension of the coefficient γ_α_ is kg/s^2−γ^, and the Laplace transform of γ(*t*) has a simple form γ_α_*s*^α-1^ (see Eq S19 in S1 Text). In the continuous limit [35], the Langevin equation of nucleosomes is described as 
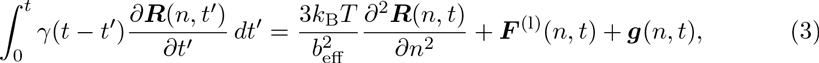
where the long-range interaction force ***F**^(l)^(n,t)* including attractive and repulsive interactions results in the size scaling 
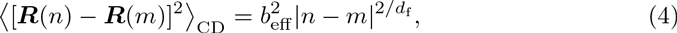

and the thermal random force ***g**(n,t)* satisfies the FDR: ⟨g_K_(n,t)g_λ_(m,t′)⟩ = *k_B_T_γ_(t — t′)δ(n − m)δ_κλ_*. Our model for *d_f_* =2 formally corresponds to the classical Rouse model in the viscoelastic medium [30], where the force ***F**^(l)^(n,t)* apparently vanishes. Hence, the additional long-range interaction force generating the scaling (Eq 4) has an important role in our model, and enables us to calculate the MSD analytically. Here, we do not take into account the hydrodynamic interactions between nucleosomes, which are discussed in Discussion and S1 Text, Section II.

### Analytical calculation shows that the MSD of nucleosomes within fractal CDs is subdiffusive in viscoelastic medium

A standard approach for treating Eq 3 is to use the normal coordinates 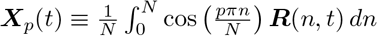 for *p* = 0,1, 2, …; however, the nonlinearity of the long-range interaction makes it difficult to deal with the equation in this manner. Therefore, to simplify the analysis, firstly, we assume that nucleosome fluctuations within the CD reach thermal equilibrium after the relaxation time *τ*_d_f__,_α_,which is explicitly described below (Eqs 11 and 12). Second, we use an approximation to transform the nonlinear Langevin equation (Eq 3) into a linear equation by averaging under thermal equilibrium with respect to the normal coordinates 
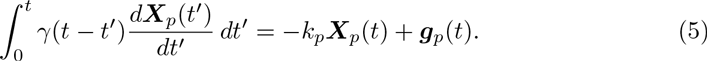

The term in the left hand side and the second term in the right hand side (RHS) are straightforwardly derived according to the normal coordinates, in which 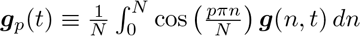 satisfies ⟨**g**_p_(t)⟩ = 0 and the FDR 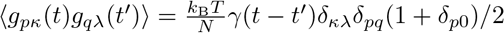 (see S1 Text, Section IA). Instead of the linearity of Eq 5, the parameter *k_P_* implicitly includes the nonlinear effect such as the long-range interactions, and is determined by the variance of ***X**_p_* over the thermal relaxation time [30] (see S1 Text, Section IB):

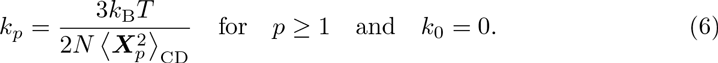

Finally, to calculate the thermal average 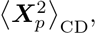, the effective size scaling (Eq 4) generated by the long-range interactions is used. The asymptotic form for large *p* is calculated as follows (see S1 Text, Section IC):

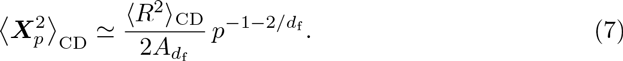

*A_d_f__* is a dimensionless constant depending on the fractal dimension:

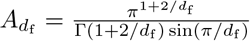. We shall refer to the above approximation as the linearization approximation, which is on the same level of the approximation as the preaveraging approximation in terms of polymer physics [35,50]. From this point forward, to avoid complicated expressions caused by this asymptotic form, we regard the asymptotic sign ‘≃’as equality.

Next, let us consider the MSD of nucleosomes in CDs. Since the inverse transform of normal coordinates is 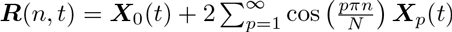 and the correlation between different modes vanishes, the MSD of the *n*-th nucleosome, ∅(*n,t*) ≡ ⟨[***R***(*n,t*) – ***R***(*n*, 0)]^2^⟩, is expressed as 
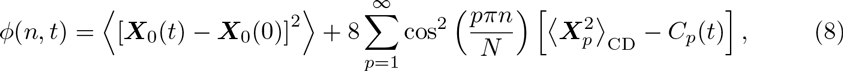
where the correlation function is defined as *C_p_*(*t*) ∅ ⟨***X**_p_*(*t*) · ***X**_p_*(0)⟩. Multiplying Eq 5 by ***X**_p_*(0) and averaging with ⟨***g**_p_*(*t*) · ***X**_p_*(0)⟩ = ⟨***g**_p_*(*t*)) · (***X**_p_*(0)⟩ = 0, we can derive that the correlation function for *p* ≥ 1 satisfies

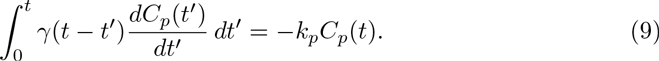

The first term for *p* = 0 in the RHS of Eq 8 corresponds to the MSD of the center of the CD, and the motion obeys 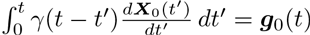 and the FDR 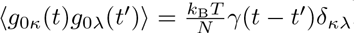. According to the fluctuation-dissipation theorem [49], the motion of the center of mass is subdiffusive with exponent α (see S1 Text, Section IE): 
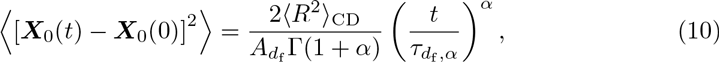
where 
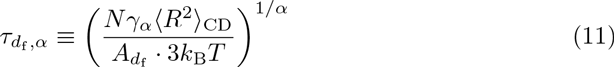
represents the relaxation time of nucleosome fluctuations in the CD.

On the other hand, the second term in the RHS of Eq 8 describes the fluctuations of many modes inside the CD. Using the Laplace transformation and the thermal equilibrium initial state, the solution of Eq 9 can be derived as follows (see S1 Text, Section ID): 
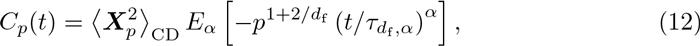
where *E_α_(x)* is the Mittag-Leffler function. According to the polymer physics [35] for *t ≪ T_d_f_,α_*, ∅(*n,t*) is dominated by terms with large *p*. Moreover, since the MSD in our experiment (Fig 2E) is calculated by averaging the nucleosome trajectories at various positions in CDs, the term 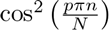 can be replaced by the average 1/2. Therefore, according to the asymptotic form of the Mittag-Leffler function, *E_α_(—x) ≃* exp [—x/Γ(1 + α)] for *x ≪* 1, and the conversion of the sum into the integral, we obtain for *t ≪ T_d_f_,α_* 
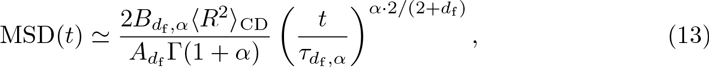
where 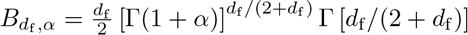 is a dimensionless constant (see S1 Text, Section IF). Thus, in our model, subdiffusive motion of single nucleosomes is a typical feature, assuming both fractal CDs and viscoelastic medium.

**Fig 2.**
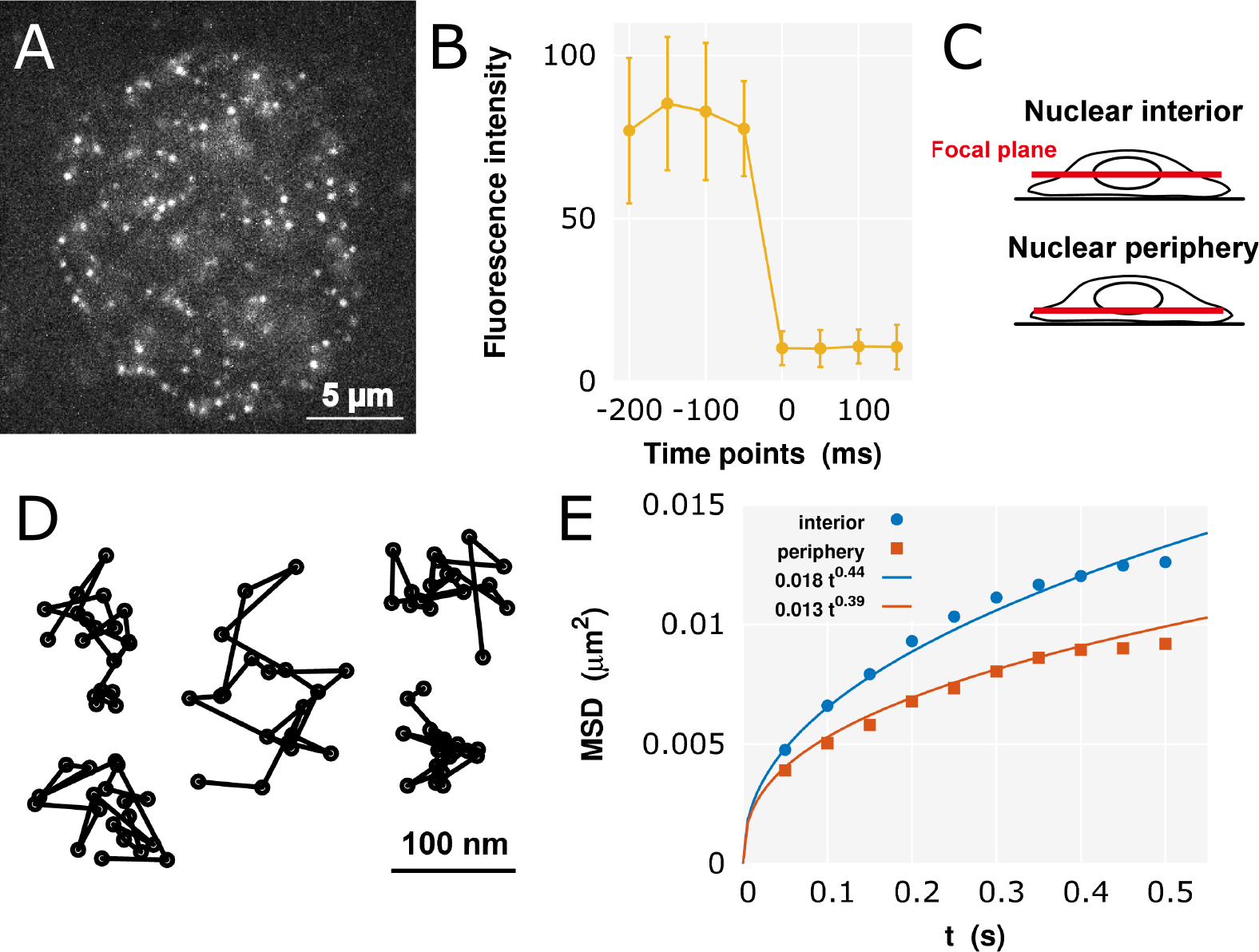
Single-nucleosome imaging and analysis. (A) Single-nucleosome image of a human HeLa cell nucleus expressing H2B-PA-mCherry. Each dot represents single nucleosome. (B) Evidence that each dot represents single-nucleosome molecule. Each H2B-PA-mCherry dot shows single-step photobleaching. The vertical axis represents the fluorescence intensity of each H2B-PA-mCherry dot. The horizontal axis is the tracking time series (each photobleaching point is set as time 0; the average and the standard deviation at each time point were calculated for 50 dots.). Due to the clear single-step photobleaching profile of the H2B-PA-mCherry, dots, each dot shows a single H2B-PA-mCherry molecule in a single nucleosome. (C) A scheme for nuclear interior (Top) and periphery (Bottom) imaging. Focal plane (red) in the living cells is shown. See also S1 Fig. (D) Representative trajectories of fluorescently labeled single nucleosome (50 ms per frame). (E) Plots of the MSD at the interior and periphery regions. These fit well with the MSD curves using Eq 1.

### Nucleosome movement is much greater in the nuclear interior than at the nuclear periphery

In order to apply our model to living human cells, single-particle imaging of nucleosomes was performed by observation of PA-mCherry labels [51] attached to histone H2B in human HeLa cells (Fig 2A). The clear single-step photobleaching profile of the H2B-PA-mCherry dots shows a single H2B-PA-mCherry molecule in a single nucleosome (Fig 2B). We tracked approximately 40,000 dots representing single nucleosomes (S1 Table). Fig 2D shows representative trajectories of the dynamic nucleosome movement in single cells.

Here, to evaluate the state of CDs according to their position in the nucleus, we focused on the nuclear interior and periphery (or surface) (Fig 2C and S1 Fig), and calculated the MSD. The nuclear periphery is a heterochromatin-rich region, which presumably shows much less active transcription than the interior. The plots of the MSD at each region, in time interval *t* up to 0.5 s, are shown in Fig 2E (normal scale) and S2 Fig (log-log scale) (also see S1 Table). The MSD at the interior is higher than that at the periphery. This result implies that nucleosome movement within CDs in the euchromatin-rich interior region is higher than that in the heterochromatin-rich periphery region.

As we analytically derived the subdiffusive MSD (Eq 13), the experimental result clearly shows subdiffusion of single-nucleosomes: using Eq 1, the plots fit well with the MSD curves 0.0181*t*^0.44^ μm^2^ and 0.0131*t*^0.39^ μm^2^ for the interior and the periphery, respectively.

### MSD is lower at the nuclear periphery than the interior, indicating that heterochromatin-rich CDs are more compact

Comparing Eqs 1 and 13, *β* and D_app_ are calculated as 
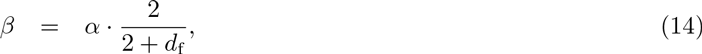

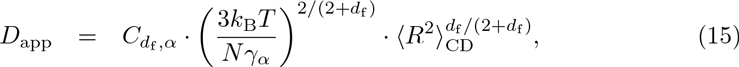
where 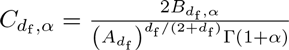. It turns out that these values contain statistical information of the CD structures, ⟨*R*⟩_CD_ and *d*_f_. Since *β* and D_app_ can be determined by the fitting in our experiments, we can therefore estimate ⟨R⟩ _CD_ and d_f_, inversely.

The lower MSD at the periphery than at the interior, D_app,periphery_ < D_app,interior_ and β_periphery_ < β_interior_, reflects the fact that the CD_s_ near the periphery are in a more compact conformational state and are smaller in size than those at the interior: *d*_f periphery_ > *d*_f,interior_ and ⟨*R*⟩_CD,periphery_ < ⟨*R*⟩_CD,interior_. This property is consistent with the conventional distribution of heterochromatin: the CDs in the heterochromatin-rich nuclear periphery are more compact than those in the euchromatin-rich interior [52].

## Discussion

To estimate the structural information of CDs through solving Eqs 14 and 15 inversely, the values of *N*, α, and *γ_α_* in mammalian living cell nuclei are required. The average size of TADs was determined to be 880 kb from mouse embryonic stem cells (mESCs), with a range of 100 kb to 5 Mb [10]. Here, we assume a CD size of 1 Mb, which corresponds to *⟨N*⟩_CD_ = 5000 nucleosomes. To the best of our knowledge, few studies have estimated the friction effect in viscoelastic cell nuclei. Therefore, we use the value of the diffusion coefficient of enhanced green fluorescent protein (EGFP)-monomer around interphase chromatin, D_EGFP_ = 20.6 μm^2^/s [25], measured by fluorescence correlation spectroscopy, in which α is assumed to be 1. In general, as a result of the FDR in a viscoelastic medium with α, the diffusion coefficient of a diffusive particle for one degree of freedom is *k_B_T /* [Γ(1 + α) · γ_α,particle_] (see Eq. S34 in S1 Text). Since the contribution of Γ(1 + α) is within the range 1 ≤ 1/Γ(1 + α) < 1.13 for 0 < α ≤1, the friction coefficient of EGFP in the nucleus can be approximately regarded as the diffusion coefficient as γ_α_→_1,EGFP_ = *k_B_T*/D_EGFP_. The hydrodynamic radius of a nucleosome bead with an H2B-PA-mCherry is assumed to be approximately quadruple for the EGFP. This means that the friction effect is also 4 times larger [48].

Accordingly, we use γ_α_→_1_ = *4k_B_T*/D_EGFP_. Finally, the structural information of CDs is estimated by calculating 
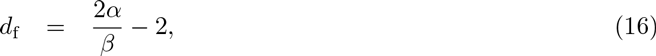

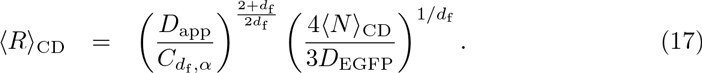
β could be measured in our experiment, although the value of α could not be determined simultaneously. Hence, Eq 16 represents the relationship between α and d_f_, as shown in Fig 3A. Under this constrained condition, according to Eqs 16 and 17, the values of the structural information within the nuclear interior and periphery regions are calculated and mapped as a function of α (Fig 3B). Since fluorescence correlation spectroscopy measurements of GFP have shown that the value of α is close to 0.79 in not HeLa but NRK nuclei [31], as an example, we summarize the estimated values for α = 0.8 and α = 0.9 in Table 1. The exponent *β* = 0.4 for the fractal globule model [28] corresponds to the value for *d*_f_ = 3 and α = 1 in Eq 14. Furthermore, our previous results have shown smaller exponents *β* = 0.37 and 0.31 for interphase chromatin and mitotic chromosome, respectively [25]. Unless considering the case of 0 < α < 1, this smaller exponent cannot be explained. Note that α has only minor effects on *C*_d_f_,α_ (see S3 Fig).

**Fig 3.**
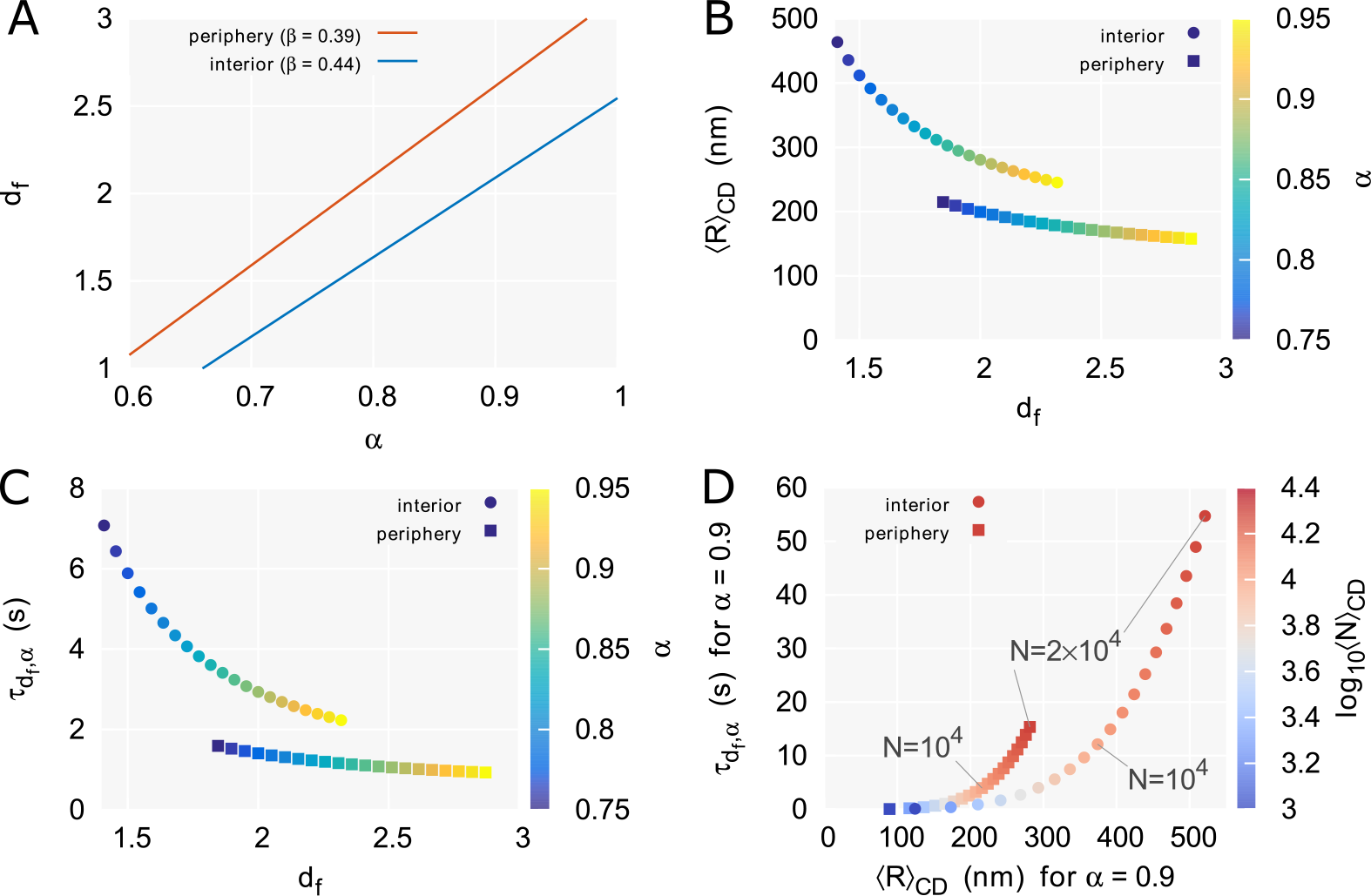
Structural information of CDs provided by single-nucleosome dynamics. (A) The fractal dimension *d*_f_ within the nuclear interior region (β = 0.44) and the periphery region (β = 0.39) for α, according to Eq 16. (B) df and the size (R) cd of, CDs, and (C) d_f_ and the relaxation time ***r**_d__f,α_* of nucleosomes in CDs within the nuclear interior region (colored circle) and the periphery region (colored square) calculated for various < values, according to Eqs 16, 17 and 18. (D) *⟨R⟩_CD_* and the *T_d__f,α_* within the nuclear interior region (colored circle) and the periphery region (colored square) calculated for α = 0.9 and various ⟨N⟩_CD_ values, corresponding to the range of 200 kb to 4 Mb, according to Eqs 17 and 18.

**Table 1.** Estimated values of the fractal dimension, d_f_, the size ⟨**R**⟩_CD_, and the relaxation time *T_d__f, α_* for α = 0.8 and α = 0.9.

| Region | $\beta$ | $D_{app}$<br>($\mu\text{m}^2/\text{s}^\beta$) | $\alpha = 0.8$ | | | $\alpha = 0.9$ | | |
| --- | --- | --- | --- | --- | --- | --- | --- | --- |
| | | | $d_f$ | $\langle R \rangle_{CD}$ (nm) | $\tau_{d_f, \alpha}$ (s) | $d_f$ | $\langle R \rangle_{CD}$ (nm) | $\tau_{d_f, \alpha}$ (s) |
| Interior | 0.44 | 0.018 | 1.64 | 358 | 4.65 | 2.09 | 268 | 2.69 |
| Periphery | 0.39 | 0.013 | 2.10 | 191 | 1.31 | 2.62 | 166 | 1.01 |

The exponent *β* and the apparent diffusion coefficient D_app_ are obtained by fitting of the MSD results at the interior and periphery regions (Fig 2C). In the, calculation, we used the following values: ⟨N⟩_CD_ = 5000 nucleosomes, *γ_α→1_* = 4kgT/D_EGFP_ and D_EGFP_ = 20.6 *μm^2^*/s.

The relaxation time of nucleosomes in CDs is calculated as 
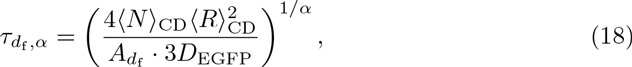
and is mapped as a function of α and d_f_ (Fig. 3C). The short relaxation time (~ s) means that the thermal equilibrium, which is the precondition for the linearization approximation, were fulfilled in our experiments. In measurements of long-term single-nucleosome movements, the MSD is expected to show a transition toward movement of the center of CDs with the exponent α (Eq 10). This would enable estimating α, d_f_, ⟨*R*⟩_CD_, and *τ_d__f_,α* without requiring the use of the assumptive values described above, such as *⟨N⟨_CD_* and D_EGFP_. The long-term (≫ *T_d__f_ α*) imaging of chromatin dynamics in mammalian nuclei might reveal this transition motion [19,21,24].

As mentioned at the beginning of this section, the measured TAD size of mESCs is in the range of 100 kb to 5 Mb. Fig 3D shows the relationship between ⟨R⟩ _CD_ and *T_d__f_,_α_* for α = 0.9 as a function of ⟨*N*⟩_CD_, corresponding to the range of 200 kb to 4 Mb, according to Eqs 17 and 18. The relaxation time within several tens of seconds is consistent with the assumption of the linearization approximation as mentioned above. Moreover, the estimated CD size within 100-500 nm is also consistent with observed radius for chromatin domains of as detected by super-resolution imaging [53].

Here, we considered a locally clustered polymer with effective size scaling (Eq 4) in the absence of hydrodynamic interactions (HIs) as a model of CDs. The inverse proportion of *k_p_* to *N*, except for the contribution from 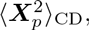 in Eq 6 reflects the lack of HIs in our model; that is, the hydrodynamic field goes through nucleosome beads without interactions. The hydrodynamic effect of surface monomers in a polymer blob on the exponent *β* has been argued in [28]. Applying their discussion to our results, Eq 16 changes into d_f_ = *c(2α/β —* 2), where the coefficient *c* is within the range 1 ≤ *c <*1.09. The effect is expected to be small. One can also consider a polymer model including HIs, which would affect the mobility matrix and work cooperatively within a polymer blob [35,50]. In such a situation, the HI cancels out the effect of the size scaling described by the fractal dimension d_f_: *β* = α · 2/3, and *β* does not depend on *d*_f_ (see S1 Text, Section II).

## Conclusion

Our results indicate that our proposed model serves as a strong method for extracting the structural information of CDs from observations of dynamic nucleosome movement. Super-resolution microscopy techniques can be used to elucidate the spatial size of CDs according to different epigenetic states [53]. On the other hand, development of an effective imaging technique to reveal the fractal dimensions remains a challenge for the future. The conformational state of CDs characterized by the fractal dimension must be associated with the accessibility of transcription factors, depending on the physical size of those factors [54]. Beyond the pioneer computational work of analyzing interphase chromosomes based on the chromatin fibers [55], further development of not only a large-scale chromosome model consisting of the CD unit based on the results of a genome-wide association study but also restraint-based three-dimensional modeling of genomes [56] is expected to provide novel insight and open the door toward further discovery on the relationship between dynamic genome organization and stochastic gene expression.

## Materials and Methods

### Cell isolation and culture

To observe single nucleosomes and analyze their local dynamics in living human cells, histone H2B was fused with photoactivatable (PA)-red fluorescent protein (mCherry) [51] and expressed in HeLa cells as described previously [25]. The cell lines expressing H2B-PA-mCherry at a very low level were isolated. The cells were cultured in Dulbecco’s modified Eagle’s medium (DMEM) supplemented with 10 % fetal bovine serum (FBS) (vol/vol) at 37°C in 5 % CO_2_ (vol/vol). The cells were plated 24-48 h before the experiment onto Iwaki glass bottom dishes treated with poly-lysine. Before the experiment, the medium was replaced by DMEM F-12 (non phenol red) with 15 % FBS. The cells were then set on the microscope stage kept in a custom-built 37°C microscope incubator enclosure with 5 % CO2 (vol/vol) delivery throughout the experiment.

### Microscopy

For single-nucleosome imaging, an oblique illumination microscope was used to illuminate a limited thin area within the cell (Nikon laser TIRF microscope system Ti with sapphire 564-nm laser). In general, PA-mCherry exhibits red fluorescence only after activation by a 405-nm laser [51]. However, we unexpectedly found that a relatively small number (~100/time frame/nucleus) of H2B-PA-mCherry molecules were continuously and stochastically activated even without UV laser stimulation. Fig 2A shows a typical single-nucleosome image of a living HeLa cell. Due to the clear single-step photobleaching profile of the H2B-PA-mCherry dots, each dot in the nucleus represents a single H2B-PA-mCherry in a single nucleosome (Fig 2B). Nucleosome signals were recorded in the interphase chromatin of the nuclear interior and periphery in living HeLa cells at a frame rate of *ca.* 50 ms/frame. Note that the two different focal planes for the nuclear interior and periphery (Fig 2C) were precisely ensured by nuclear surface labeling with Nup107 (a nuclear pore component)-Venus (a bright yellow fluorescent protein) [57] (see S1 Fig).

### Tracking and data analysis

Local nucleosome fluctuation was observed (*ca*. 60 nm movement/50 ms), presumably caused by Brownian motion. The free MATLAB software u-track [58] was used for single-nucleosome tracking. The dots were fitted to an assumed Gaussian point spread function to determine the precise center of the signals with higher resolution. Finally, we obtained data set of two-dimensional *M_i_* trajectories 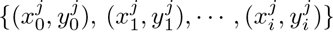, where the suffix *j ∊* {1, …, *M_i_α* represents the sample number for the tracked time-interval [0,t_i_]; *t_i_ ≡ i* × 50 ms. Several representative trajectories of fluorescently tagged single nucleosomes are shown in Fig 2D (bar = 100 nm).

According to observed regions, we calculated the ensemble-averaged MSD of single nucleosomes: 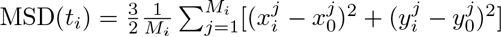. Here, in order to obtain the three-dimensional value, we multiplied the two-dimensional value by 3/2 on the assumption of isotropy. Plots of the MSDs of single nucleosomes in interphase chromatin at the nuclear interior (10 cells) and the nuclear periphery (10 cells) from 0 to 0.5 s are shown in Fig 2E. The plots for single nucleosomes were fitted with the subdiffusion model (Eq 1) using R-software. The standard error of the mean (SEM), which is the standard deviation of the sampling distribution of the mean, for MSD(t_i_) was sufficiently small. The number of trajectories *M_i_* and the SEM of MSD(t_i_) are summarized in S1 Table.

## Acknowledgments

The authors thank Shin-ichi Tate, Akinori Awazu, Takeshi Sugawara, Hiroshi Ochiai, and Yasunori Horikoshi for fruitful discussions.

## Supporting Information

S1 Text. Further details on derivations of the theoretical results and remarks on the hydrodynamic effect for the model.

**S1 Fig.**
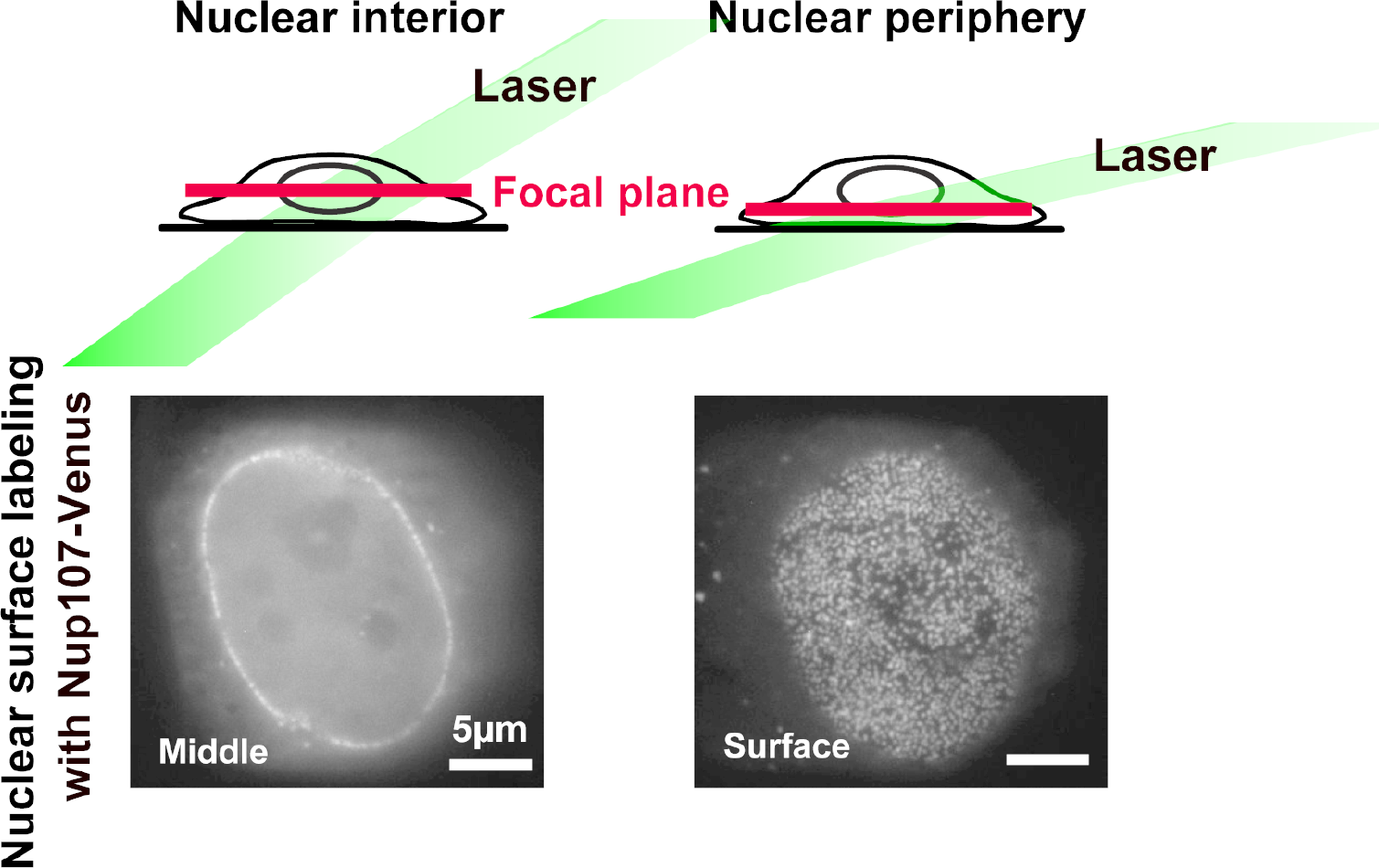
A schematic representation for nuclear interior (Top left) and periphery (Top right) imaging. Illumination laser (green) and focal plane (red) in the living cells are shown. Note that the two different focal planes were precisely verified by nuclear surface labeling with Nup107 (a nuclear pore component)-Venus (a bright yellow fluorescent protein) [57]. The nuclear rim signals (Bottom left) and dot signals in ellipse shape (Bottom right) show the middle layer of nucleoplasm and the nuclear, surface, respectively. Bar shows 5 μm.0.1

**S2 Fig.**
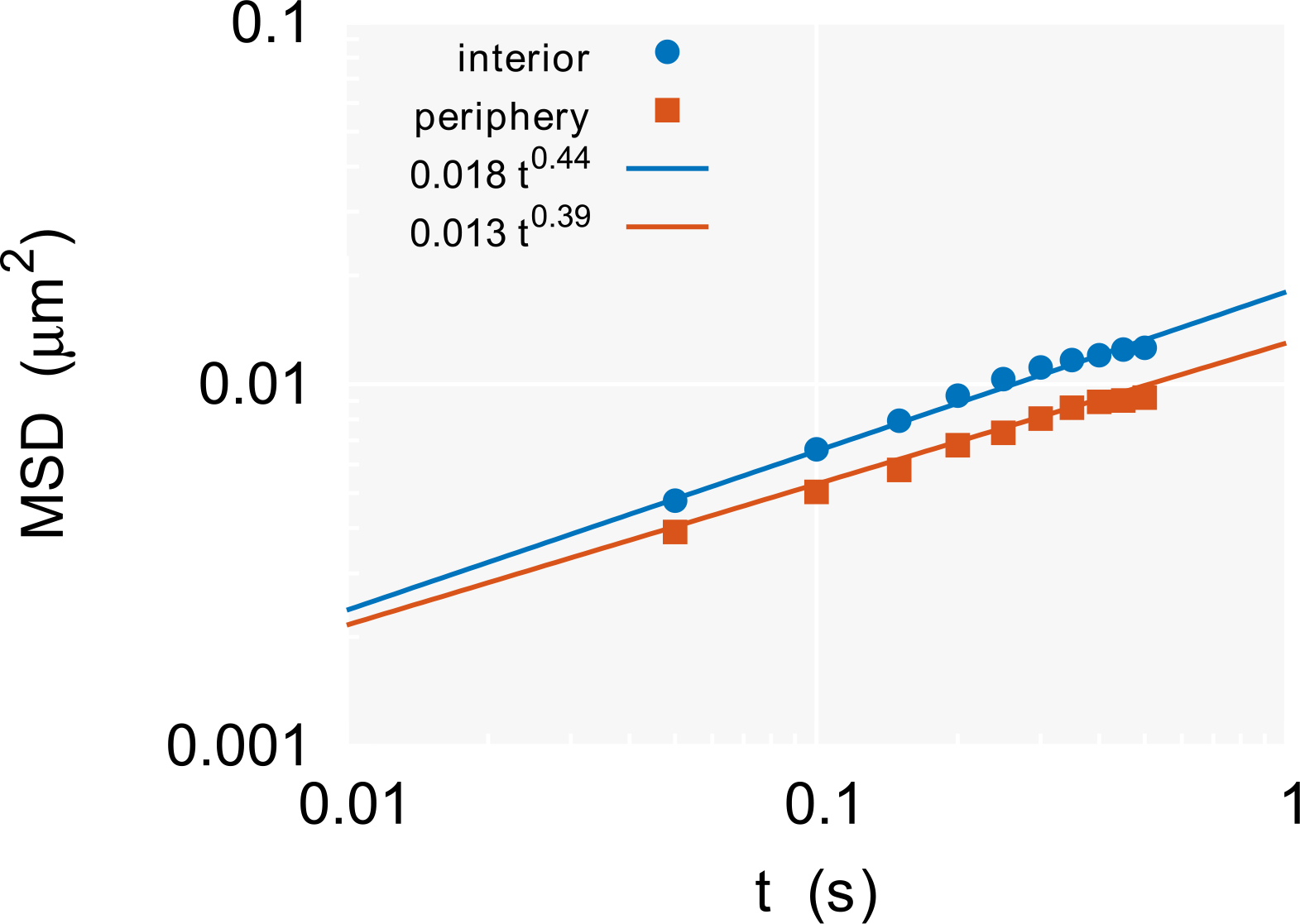
Plots of the MSD (Fig 1E) on log-log scale.

**S3 Fig.**
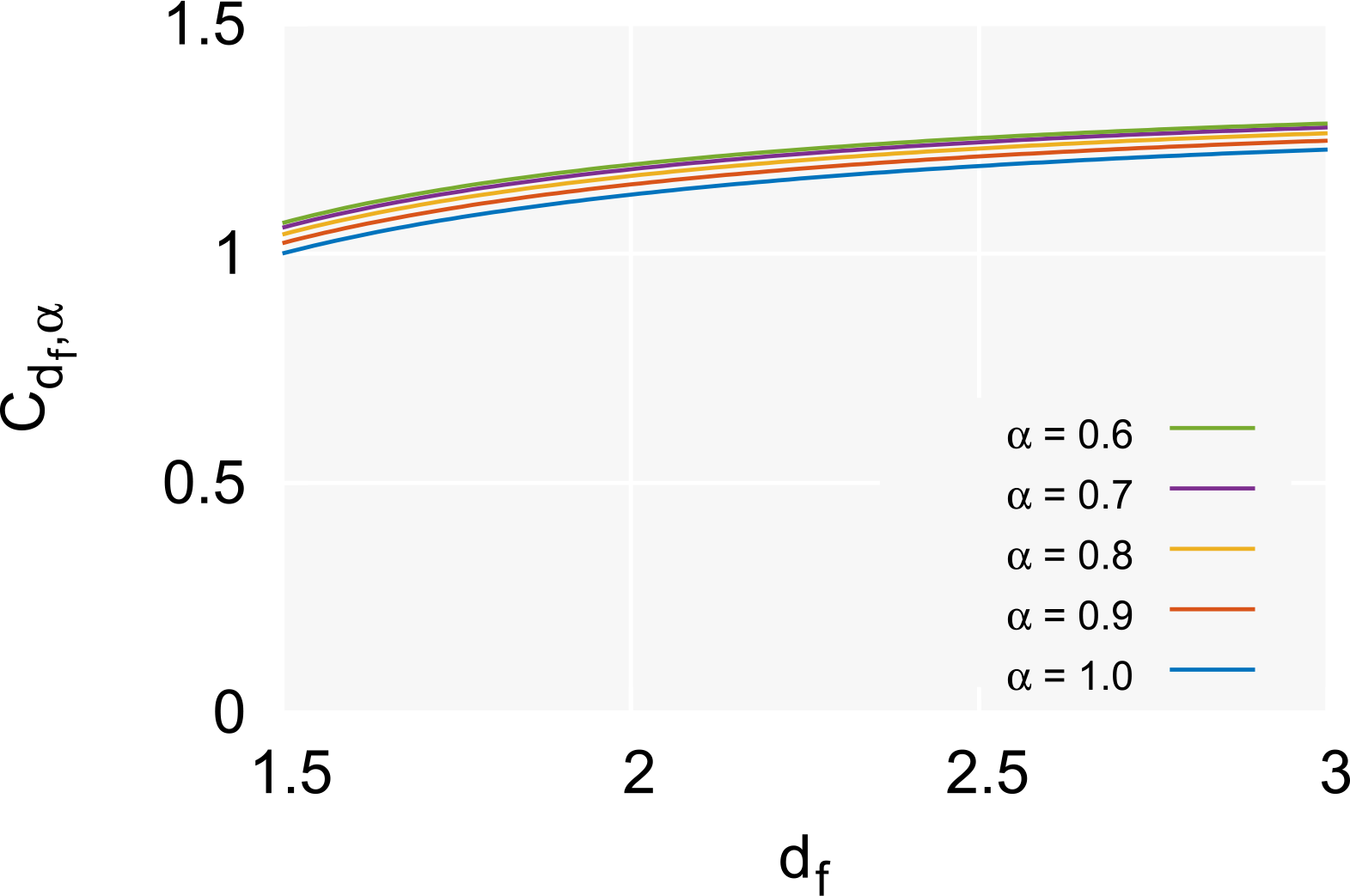
The function C_d__f,α_ of the fractal dimension d_f_ for α = 0.6, 0.7, 0.8, 0.9, and 1.0. α has only a slight effect on C_d__f,α_.

**S1 Table.** **The number of tracked trajectories *M_i_* and the standard error of the mean (SEM) of the MSD at the nuclear interior region and the periphery region**. The measurements at each region were performed using 10 cells.

| $t_i$ (ms) | Interior | | Periphery | |
| --- | --- | --- | --- | --- |
| | $M_i$ | SEM of MSD($t_i$) | $M_i$ | SEM of MSD($t_i$) |
| 50 | 47797 | 0.0000246 | 35289 | 0.0000264 |
| 100 | 47732 | 0.0000361 | 35267 | 0.0000367 |
| 150 | 47498 | 0.0000424 | 35176 | 0.0000418 |
| 200 | 36291 | 0.0000550 | 26072 | 0.0000556 |
| 250 | 27876 | 0.0000683 | 19614 | 0.0000686 |
| 300 | 21625 | 0.0000817 | 14572 | 0.0000839 |
| 350 | 17074 | 0.0000957 | 11008 | 0.0001017 |
| 400 | 13650 | 0.0001093 | 8533 | 0.0001190 |
| 450 | 11047 | 0.0001240 | 6825 | 0.0001331 |
| 500 | 9047 | 0.0001371 | 5491 | 0.0001510 |

## I. DERIVATIONS OF THE THEORETICAL RESULTS

Here we give the complete derivations of the theoretical results. The following calculations are based on standard textbooks related to polymer dynamics [S1] and statistical physics [S2]. The mean-squared displacement (MSD) of the Rouse polymer in the viscoelastic environment was first analyzed by Weber *et al.* [S3], whose work is a useful reference for the flow in the following calculations.

### A. The fluctuation-dissipation relation between *g_p_(t)* and *γ(t)*

According to the fluctuation-dissipation relation (FDR) for ***g**(n,t),* 
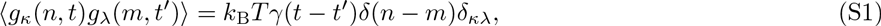
the following calculations can be made:
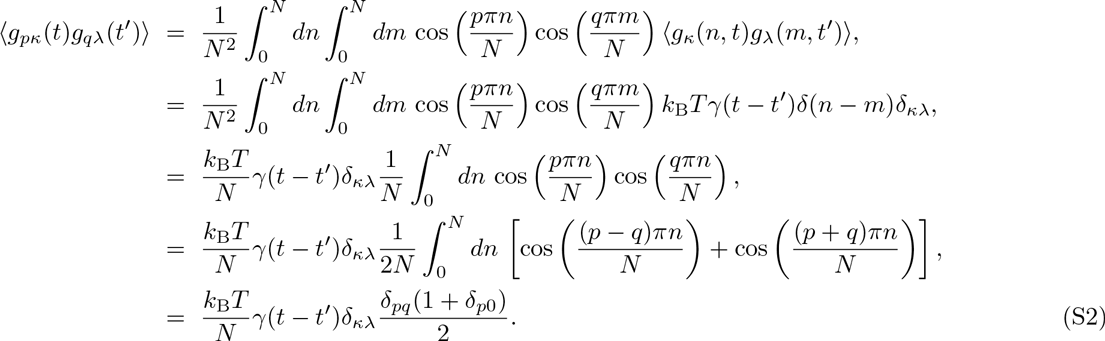

### B. The parameter *k_p_* relates to the variance of *X_p_*

At thermal equilibrium via the preaveraging, approximation, the memory effect of the friction coefficient, vanishes, *i.e.,* 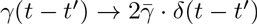. Then, Eq. 5 for *p* ≥1 can be written as 
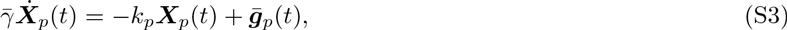
where 
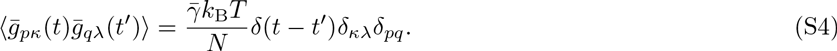

Since this Langevin equation for one degree of freedom corresponds to the Ornstein-Uhlenbeck process described by the stochastic differential equation [S4], 
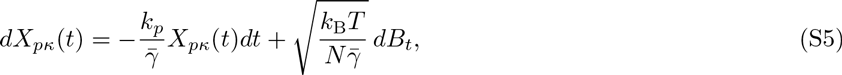
the variance of *X_p_* becomes 
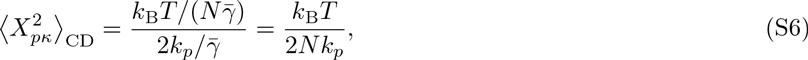
where ⟨·⟩_CD_ represents the average for all nucleosome beads within the chromatin domain (CD) at thermal equilibrium. Thus, this relation implies that the normal-coordinate amplitude satisfies the equipartition theorem at thermal equilibrium.

### C. Asymptotic form of 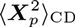

Here, we omit the argument *t* to calculate the thermal average. Using integration by, parts, the normal coordinates 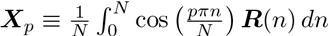 
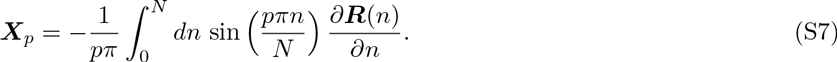

Thus, 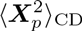 is written as 
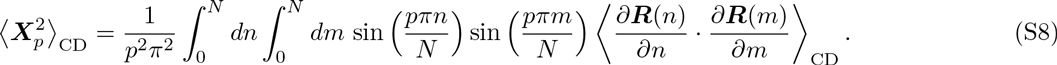

Using 
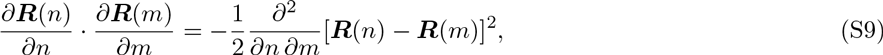
we can rewrite 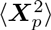 as 
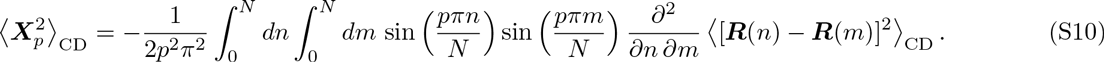

Introducing a new variable *l* = *m* — *n* and substituting the size scaling (Eq. 4), we can make the following calculation: 
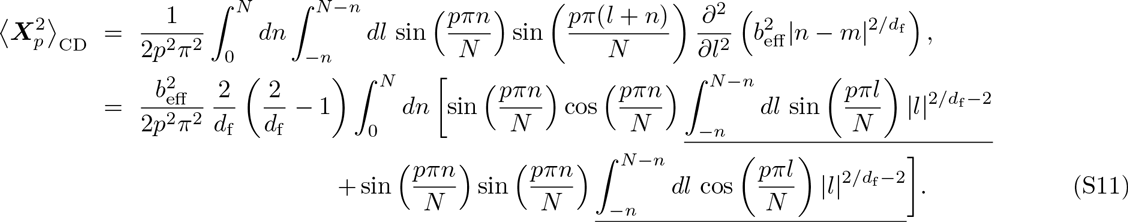

The underlined integrals converge quickly to the following values if *p* is large: 
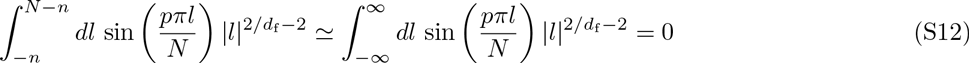
and 
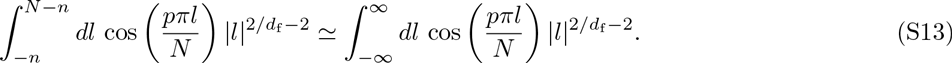

Therefore, we can obtain 
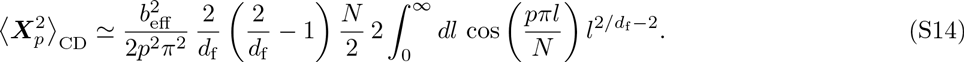

Using the formulas 
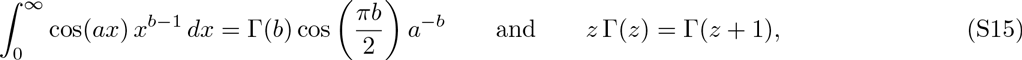
we can make the following formal calculations: 
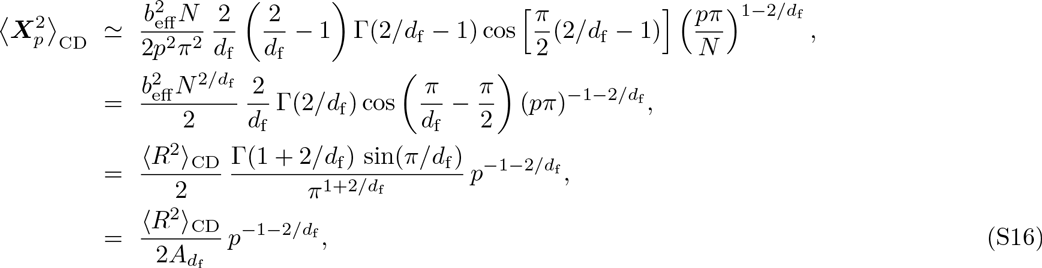
where 
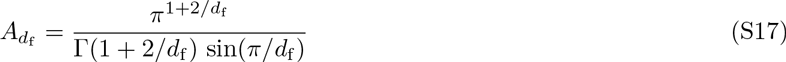
is a dimensionless constant depending on the fractal dimension *d_f_*.

### D. The solution of Eq. 9

Performing the Laplace transform to Eq. 9, we obtain 
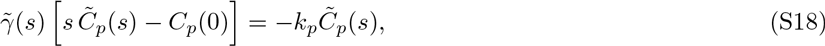
where 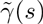 and 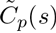 are the Laplace transforms of the functions *γ*(*t*) and *C_p_*(*t*), respectively. Since *γ*(*t*) is defined by Eq. 2, 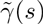 is derived as follows: 
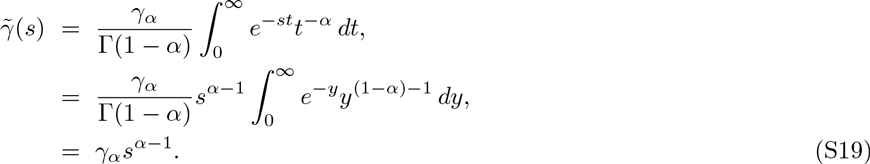

Therefore, 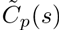 is written as 
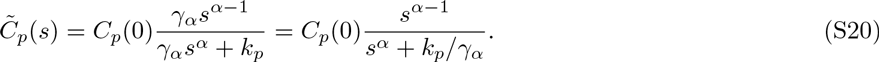

In, addition, using the formula of the Laplace transform for the Mittag-Leffler function 
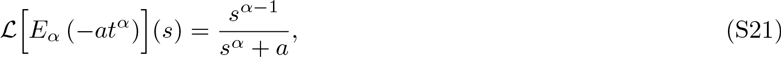
we can inversely find the solution 
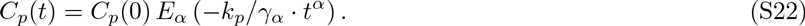

By use of Eqs. 6 and 7, 
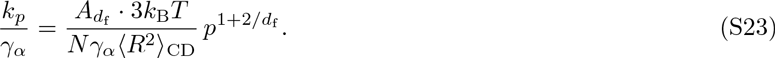

Then, we can define the relaxation time 
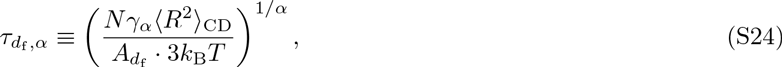
which has the physical dimension s. If the initial condition reaches thermal, equilibrium, *C_p_*(0) becomes 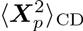. Thus, finally, we can derive the solution 
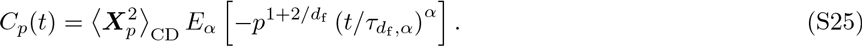

### E. The MSD of the center of the CD

For *p* = 0, the normal coordinate **X**_0_(t) corresponds to the center of the, CD, 
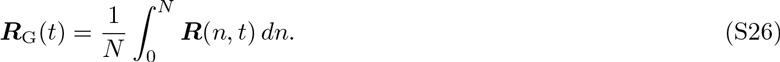

According to the Langevin equation (Eq. 5) and the FDR (Eq. S2) for *p* = 0, the motion obeys 
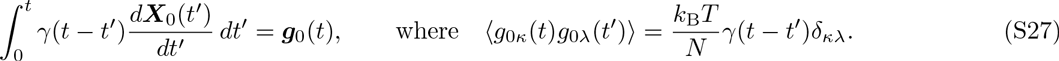

In, general, for degree of freedom *x* and velocity *v*, the MSD is associated with the velocity correlation as follows: 
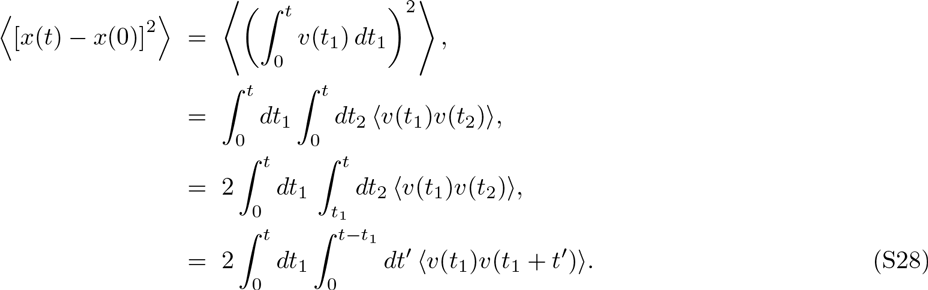

Using the Laplace transform and the stationarity of the velocity correlation *C_v_*(t), this relation becomes more clear: 
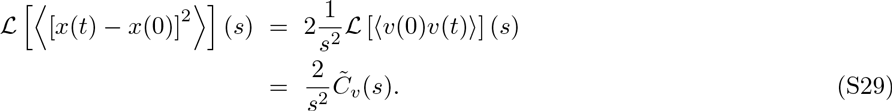

In terms of the fluctuation-dissipation theorem (FDT) [S2], we can derive the Laplace transform of the velocity correlation from the relationship between the average response and the FDR. The force balance between the average response of the system described by Eq. S27 and the external force *f*(*t*) is written as 
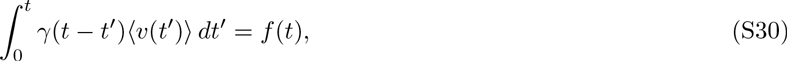
for one degree of freedom. The Laplace transform of this force balance equation becomes 
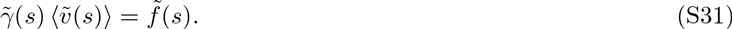

Then, the ratio of the average velocity to the force, 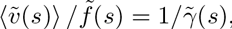, is called the complex admittance, and the FDT of the first kind represents the relationship between the complex admittance and the velocity correlation, 
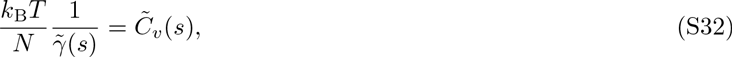
where the coefficient kBT=N is caused by the FDT of the second kind for the system in Eq. S27.

Therefore, the Laplace transform of the MSD is written as 
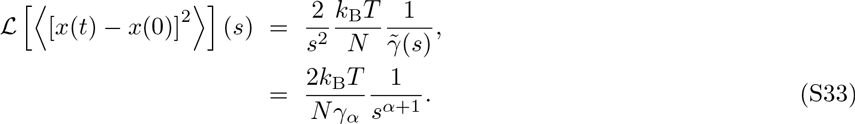

By use of the formula of the inverse Laplace transform, L^−^1[1/s^α+1^](t) = t^α^/Γ(1 + α),the MSD can be obtained as 
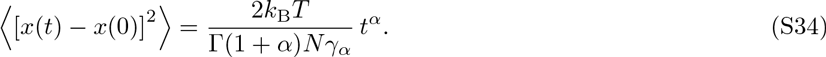

Thus, the MSD of the center of the CD is derived as 
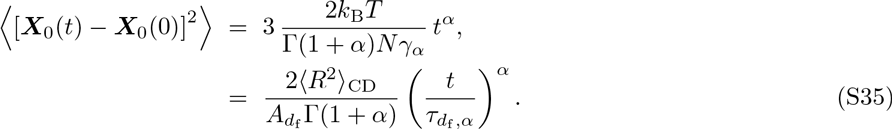

### F. The MSD for t ≪ t_d_f_,α_

The MSD obtained in our experiment is calculated by averaging nucleosome movements at various positions in CDs. Then, we can replace the term 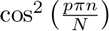 in Eq. 8 by the average 1/2. Therefore, for t ≪ *τ* _d__f,α_, according to Eqs. 7 and 12, and the asymptotic form of the Mittag-Leffler function, E_α_(—x)≃ exp[— x/Γ(1 + α)] for x ≪ 1, the second term in the right hand side (RHS) of Eq. 8 can be expressed as 
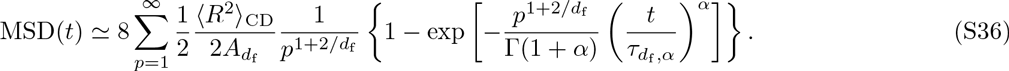

Converting the sum into the integral, the RHS becomes 
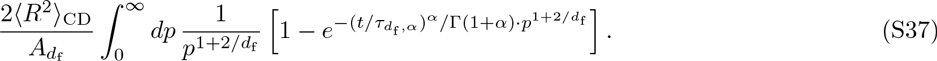

Here, let us consider the integral formula calculated as follows: 
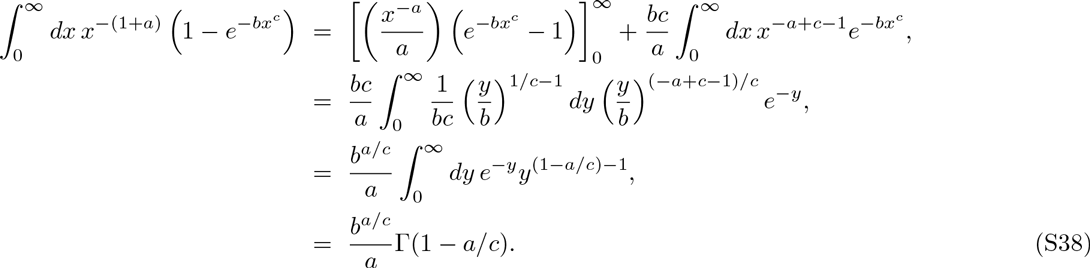

Therefore, the MSD for t ≪ T_d_f_,α_ can be written as 
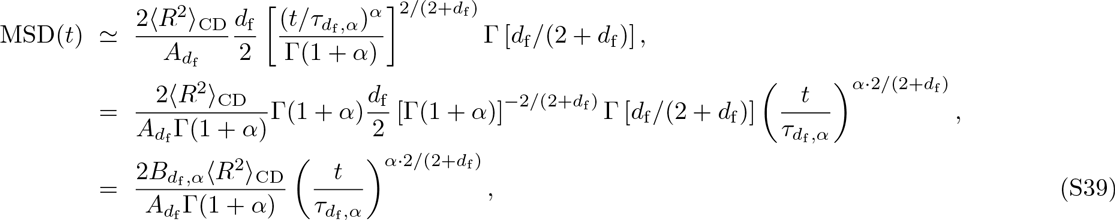
where 
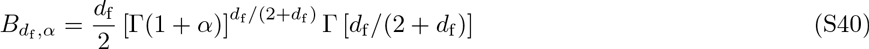
is a dimensionless constant depending on d_f_ and α.

## II. REMARKS ON THE HYDRODYNAMIC EFFECT FOR OUR POLYMER MODEL

In describing the Langevin equation of polymers with the hydrodynamic interaction, the interaction affects the mobility matrix [S1, S5]. This situation corresponds to an ideal case where hydrodynamic interactions are not screened. Calculating the effect of the mobility matrix for the normal coordinates **X**_p_(t) under the preaveraging approximation, *k_p_* in Eq. 5 is changed into 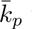with the following *p*-dependence: 
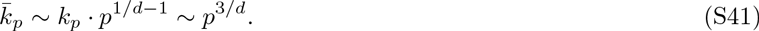

Therefore, when we calculate the MSD as above, we need to calculate the integral 
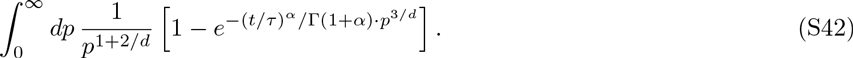

By use of the integral formula (Eq. S38), the scaling of the MSD for t ≪ *τ* can be written as 
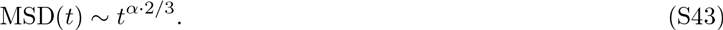

This means that the hydrodynamic interaction cancels out the effect of the size scaling described by the fractal dimension d_f_, and that the exponent of the MSD depends on only the exponent α, which relates to the memory effect of the viscoelastic medium.

